# The Nucleoporin Nup153 Is the Anchor for Kif1a during Basal Nuclear migration in Brain Progenitor Cells

**DOI:** 10.1101/2023.07.25.550108

**Authors:** Aditi Falnikar, Sebastian Quintremil, Paige Helmer, Richard B. Vallee

**Affiliations:** Department of Pathology and Cell Biology, Columbia University Medical Center, New York, NY, USA

**Author notes:** Co-corresponding authors: Richard B. Vallee, Aditi Falnikar.

**Keywords:** Kif1a, Nup153, brain development, interkinetic nuclear migration, neuronal migration, nuclear transport

## Abstract

Radial glial progenitors (RGPs) are highly elongated epithelial cells that give rise to most stem cells, neurons, and glia, in the vertebrate cerebral cortex. During development the RGP nuclei exhibit a striking pattern of cell cycle-dependent oscillatory movements known as interkinetic nuclear migration (INM), which we previously found to be mediated during G1 by the kinesin Kif1a, and during G2 by cytoplasmic dynein, recruited to the nuclear envelope by the nucleoporins RanBP2 and Nup133. We now identify Nup153 as a nucleoporin anchor for Kif1a, responsible for G1-specific basal nuclear migration, providing a complete model for the mechanisms underlying this basic, but mysterious behavior, with broad implications for understanding brain development.

## MAIN TEXT

RGP nuclei undergo a series of cell cycle-dependent oscillatory movements known as interkinetic nuclear migration (INM)^1-3^. Mitosis begins when the G2 nucleus reaches the apical end of the cell at the ventricular surface (VS) of the brain. The nucleus then migrates basally during G1, completes S-phase away from the ventricle and returns to the VS during G2, for the next mitotic division. Symmetric and asymmetric RGP divisions respectively increase progenitor numbers or yield new neurons, which migrate to the cortical plate (CP). INM may be involved in cell fate determination^4^ or in maximizing the number of dividing cell bodies accommodated at the VS^5^.

We have reported roles for cytoplasmic dynein and Kif1a in INM and post-mitotic neuronal migration using fixed and live imaging of embryonic rat brain^2, 3, 6, 7^. The centrosomes remained at the VS during INM, with microtubules (MTs) within the apical processes, oriented uniformly with their minus ends toward the pial brain surface. Consistent with this arrangement, RNAi for the MT plus end-directed kinesin, Kif1a, inhibited basal nuclear migration, whereas RNAi for cytoplasmic dynein or its regulator LIS1 inhibited apical migration.

Although centrosomes associate with nuclei during migration in many cell types, the centrosome-independent nuclear migration we observed in RGPs^3, 8^ suggested that the motor proteins in this case might act locally at the nuclear surface. In the case of rat RGPs, we found sequential G2 mechanisms responsible for dynein nuclear pore recruitment, involving, respectively, RanBP2-BicD2-dynein and Nup133-CENP-F-dynein interactions^2^.

In contrast to dynein recruitment to the RGP nuclear envelope, nothing is known of an equivalent mechanism for Kif1a. Kif1a has well-established roles in higher eukaryotic transport of synaptic vesicle precursors^9^ and other vesicular cargo. Human KIF1A mutations have also been implicated in severe, progressive neurological disease^10, 11^.

To identify potential NE-recruitment factors for Kif1a we took advantage of data from a recent interactome study^12^, and focused on four Kif1a-interacting nucleoporins, Nup153, Nup88, Nup160, and Nup214. Of these, Nup153 alone had a high likelihood of interaction and was therefore tested for its role in INM. We used Nup160 as a negative control in these experiments.

To test for roles in INM, we expressed shRNAs corresponding to each Nup in E16 rat brain (Supplemental Figure 1). Nup153 knockdown caused significant accumulation of RGP somata within 10 μm of the VS though cell morphology was unaffected, and basal processes still extended to the pial surface of the brain. Neither Nup160 nor scrambled Nup153 shRNAs affected the distribution of RGPs somata (Figure 1).

**Figure 1.**
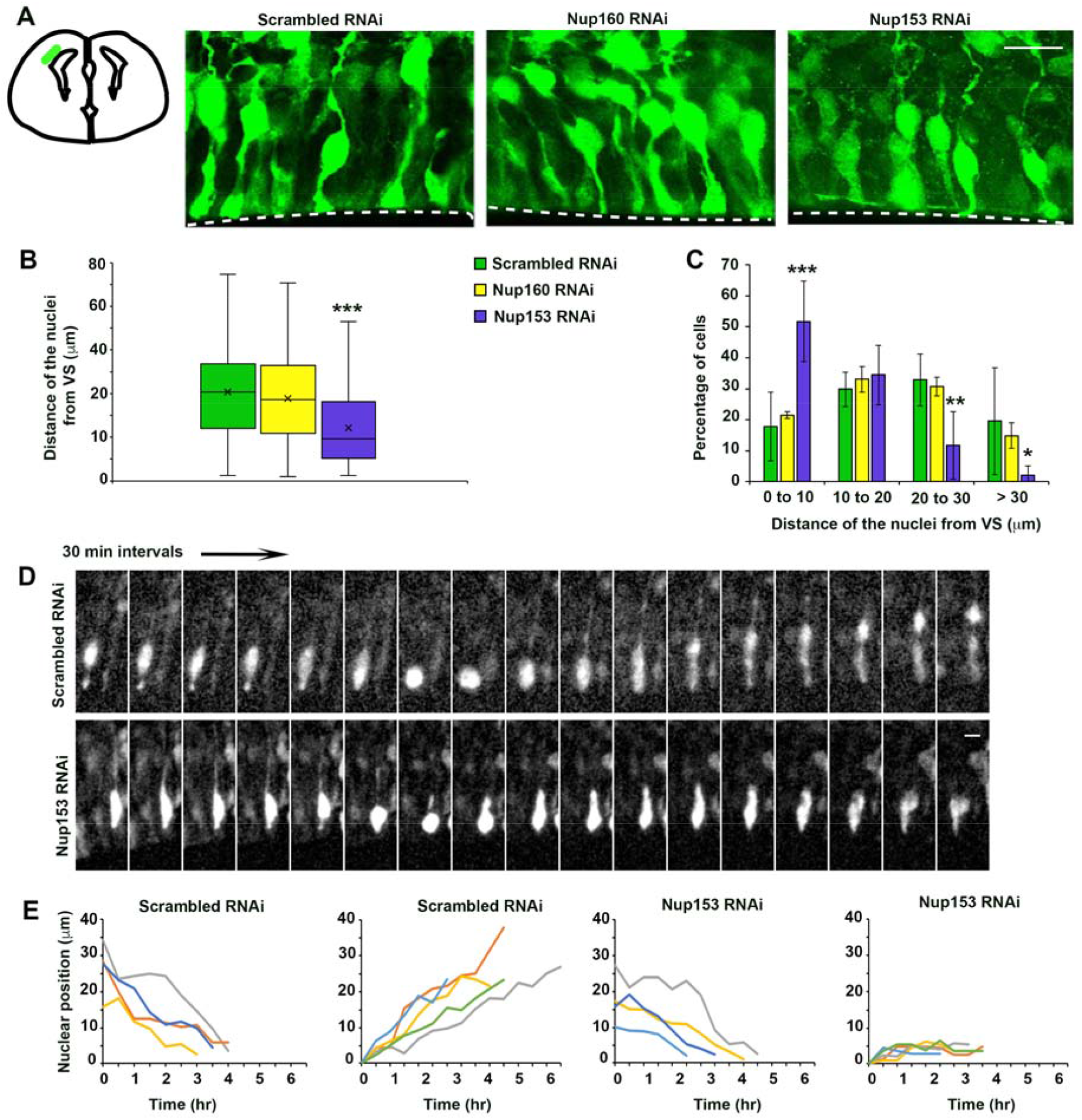
Nup153 is required for basally directed nuclear movement in radial glial progenitors. E16 embryonic brains were in utero electroporated with scrambled shRNA or shRNAs for Nup160 or Nup153 and subsequently imaged live or fixed 3 days post injection, at E19. **A**. Fixed images of the VZ from electroporated brains stained for EGFP. Dashed line represents the VS. KIF1A RNAi caused clear accumulation of RGP somata near VS. Presence of radial glial basal process indicates retention of overall cell morphology. Scale bar 15 μm. **B and C**. Quantification of distance between the RGP nuclei and the VS across conditions. Control nuclei were distributed at a range of distances from the ventricular surface. Nuclei of Nup153 shRNA expressing cells accumulated relatively closer to the VS. (0 – 10 μm: scramble shRNA, 17.8 ± 4.2%, n = 7; Nup160 shRNA, 21.5 ± 0.6%, n = 3, p = 0.8; Nup153 shRNA, 51.7 ± 5.3%, n = 6; p = 0.0004; 10 – 20 μm: scramble shRNA, 29.8 ± 2.1%, n = 7; Nup160 shRNA, 33.0 ± 4.1%, n = 3, p = 0.6; Nup153 shRNA, 34.5 ± 3.9%, n = 6, p = 0.3; 20 – 30 μm: scramble shRNA, 32.8 ± 3.2%, n = 7; Nup160 shRNA, 30.7 ± 1.8%, n = 3, p = 0.6; Nup153 shRNA, 11.7 ± 4.5%, n = 6, p = 0.002; >30 μm: scramble shRNA, 19.5 ± 6.5%, n = 7; Nup160 shRNA, 14.8 ± 2.4%, n = 3, p = 1; Nup153 shRNA, 2 ± 1.9%, n = 6, p = 0.03; _∗∗_p < 0.01; _∗∗∗_p < 0.001; error bars = SEM) **D**. Time-lapse images for scrambled shRNA or Nup153 shRNA in RGPs (Videos 1 and 2). Images are shown at 30-min intervals. Scale bar 10 μm. **E**. Tracings of movements of individual nuclei in scrambled or Nup153 shRNA-expressing RGP cells. There was no apparent effect of Nup153 RNAi in apical direction. In contrast, marked effects are observed in basal direction for all RGP nuclei.

To test the role of Nup153 more directly, we performed live imaging of RGPs. Consistent with fixed tissue analysis, live cell imaging revealed specific inhibition of basal INM in RGPs, with nuclei remaining within ∼15 μm of the VS throughout the recording period (Figure 1). Some short departures from the VS could be seen, possibly due to residual Nup153 expression or passive nuclear displacement. These results together, suggest that Nup153 RNAi phenocopies Kif1a RNAi.

We previously found that Kif1a RNAi does not affect cell cycle progression^6^, suggesting that cell cycle progresses normally despite inhibition of basal INM. To test for the effect of Nup153 RNAi, we stained Nup153 shRNA expressing brain sections with cell cycle markers. While Nup153 RNAi had no gross effect on the fraction of Ki67-positive cycling cells, it caused a significant increase in Cyclin D1-positive G1 phase cells with a concomitant decrease in cells in S-phase as well as in G2/M-phase (Supplemental Figure 2).

To understand the relative role of Nup153’s functional domains in anchoring Kif1a to the RGP nucleus, we next expressed previously characterized cDNA’s encoding individual Nup153 domains^13^ in E16 brains. The N-terminal portion of Nup153 (Nup153-N) contains the nuclear envelope (NE)- and nuclear pore complex (NPC)- targeting sequences^14, 15^ and localizes to NE in HeLa cells. The Nup153 C-terminal domain (Nup153-C) is flexible and extends to the cytoplasmic surface of NPC^16-18^. This domain can be detected in both the cytoplasm and nucleus when expressed in HeLa cells (Supplemental Figure 4). Expression of Nup153-C or Nup153-full length constructs in E16 brains had no detectable effect on RGP nuclear distribution. Interestingly, expression of Nup153-N resulted in a pronounced impairment of basal INM, similar to Nup153 KD (Supplemental Figure 3). Together, these results suggest the Nup153-N competes with endogenous Nup153 for NPC incorporation but is, by itself, insufficient to anchor Kif1a to the nucleus. We reason that this is because the Kif1a-binding site lies elsewhere within the Nup153 molecule. These results confirm the role of Nup153 in anchoring Kif1a to the nucleus for basal nuclear migration.

To further understand the role of Nup153 in NE Kif1a recruitment we tested for co-distribution of Nup153 with endogenous as well as recombinant Kif1a in nonneuronal cells. Interestingly, Nup153-C showed a striking pattern of co-localization with the endogenous Kif1a puncta (Supplemental Figure 4).

In addition to its microtubule-binding motor domain, Kif1a has a large C-terminal “stalk” (hereafter “tail”) which contains binding sites for several known forms of membranous cargo. This tail region of Kif1a (amino acid 657-1105) was originally used to pull down Nup153 from adult rat brain lysate^12^. When expressed by itself, Kif1a tail domain showed a clear NE decoration in a majority of HeLa cells (Figure 2), consistent with an interaction with Nup153. When co-expressed with Nup153, we again observed extensive diffuse as well as punctate colocalization of Nup153-C with the Kif1a tail region (Supplemental Figure 5).

**Figure 2.**
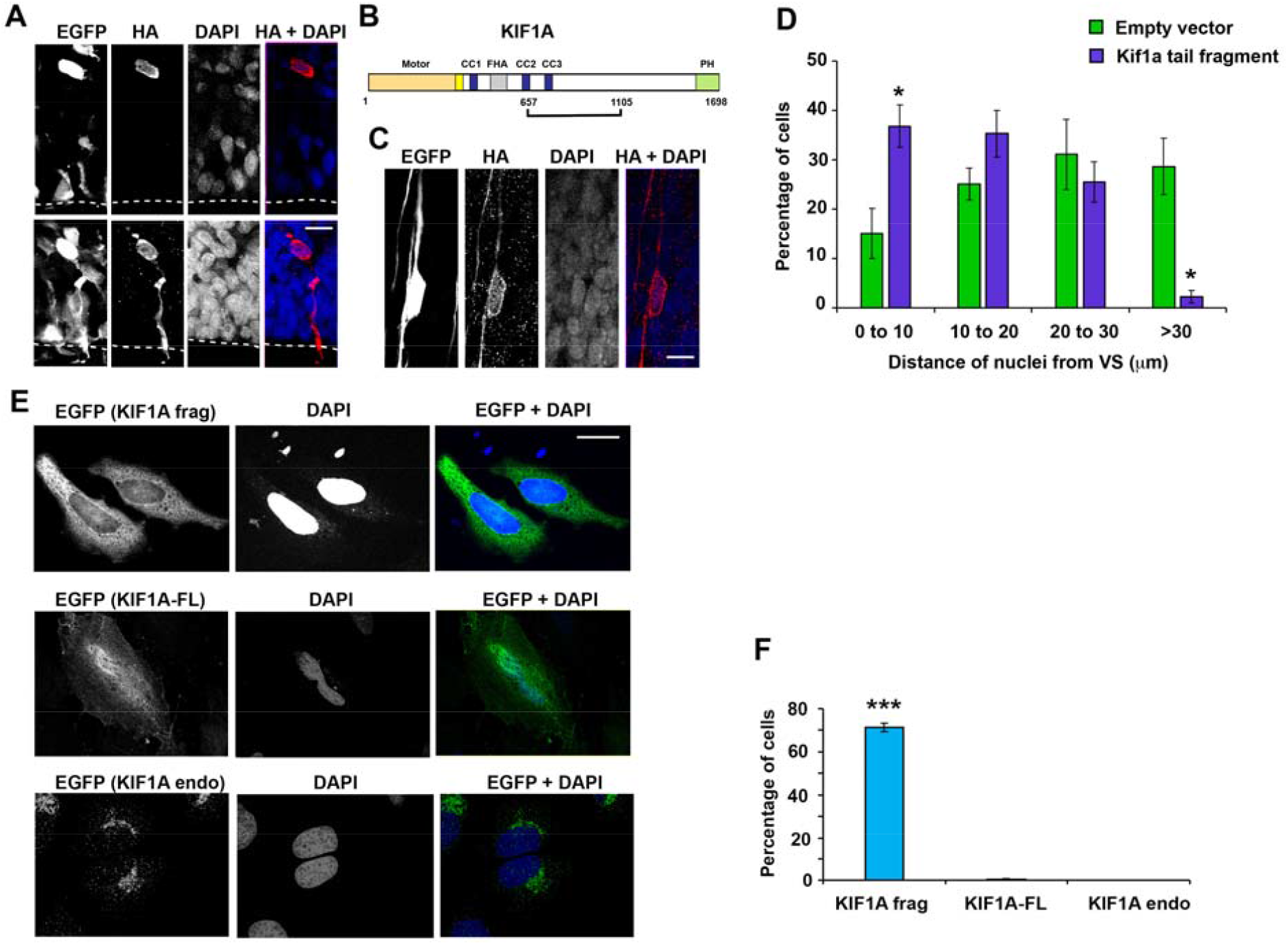
KIF1A tail fragment decorates nuclear envelope (NE) in RGP cells, migrating neurons and HeLa cells and its expression in RGP cells inhibits basal INM. **A, C**. E16 embryonic brains were in utero electroporated with the KIF1A tail fragment with HA-tag that is fused at its N-terminal and EGFP-tag that is cytoplasmic; and subsequently imaged following fixation 3 days post injection, at E19. **A**. Fixed images of the ventricular zone from the electroporated brains stained for EGFP and HA. Dashed lines represent the VS. HA staining showed cleared NE decoration in a subset of RGP cells. Scale bar 10 μm. **C**. Fixed images of the cortical plate region from the electroporated brains stained for EGFP and HA. HA staining showed cleared NE decoration in a subset of migrating neurons. Scale bar 10 μm. **B**. Schematic representation of the KIF1A molecule and its tail fragment that was used in this study. **D**. Quantification of the fixed images of the VZ from electroporated brains expressing either KIF1A tail fragment or control empty vector, representing the distance between the RGP nuclei and the VS between the two conditions. Control nuclei were distributed at a range of distances from the ventricular surface. Nuclei of KIF1A tail fragment expressing cells accumulated relatively closer to the VS (0 – 10 μm: control, 15.1 ± 5.1%, n = 3; Kif1a tail, 36.8 ± 4.3%, n = 3, p = 0.03; 10 – 20 μm: control, 25.1 ± 3.2%, n = 3; Kif1a tail, 35.3 ± 4.7%, n = 3, p = 0.1; 20 – 30 μm: control, 31.1 ± 7.1%, n = 3; Kif1a tail, 25.5 ± 4.1%, n = 3, p = 0.5; >30 μm: control, 28.9 ± 5.7%, n = 3; Kif1a tail, 2.3 ± 1.3%, n = 3, p = 0.01; _∗∗_p < 0.01; _∗∗∗_p < 0.001; error bars = SEM). **E**. HeLa cells expressing KIF1A tail fragment with EGFP-tag that is fused at its N-terminal, fixed and stained for EGFP. EGFP staining shows clear NE decoration in a subset of cells. Whereas, NE decoration was not observed in HeLa cells expressing either full length KIF1A or the ones stained with the anti-KIF1A antibody to detect endogenous KIF1A. Scale bar 20 μm. **F**. Quantification of the fixed images of the HeLa cells expressing either KIF1A tail fragment or full length KIF1A and stained for EGFP to detect expression at the NE. Also included is the third condition where HeLa cells were stained with anti-KIF1A antibody to detect the endogenous Kif1a expression at the NE. Only the HeLa cells expressing KIF1A tail fragment showed NE localization. Each experiment was reproduced three independent times (over 50 cells per condition and per experiment were counted. _∗∗_p < 0.01; _∗∗∗_p < 0.001; error bars = SEM).

Multiple proteins, including Nup153, were found associated with the KIF1A tail region in an earlier study^12^. The effect of Nup153 KD on basal INM supports our original proposal for Kif1a-mediated forces generated at the NE. To test whether the Kif1a tail region affects basal INM, we electroporated a cDNA encoding EGFP-as well as HA-tagged Kif1a tail region into E16 rat brain. As in HeLa cells, we observed clear NE decoration in most of the transfected RGP cells as well as in postmitotic neurons (Figure 2). Expression of this domain also inhibited basal INM (Figure 2) suggesting that free Kif1a tail domain might compete with endogenous Kif1a for a common binding site at the NE. Our results, together, are consistent with the role of Nup153 in anchoring Kif1a, via its C-terminal domain, to the NE to drive basal nuclear migration.

Previous work from our lab and others suggested Kif1a participates in post-mitotic neuronal migration as well^6, 19^. Little is known about the mechanisms responsible for this behavior. To test whether this behavior shares mechanistic features with INM, we performed Nup153 RNAi and looked at the behavior of postmitotic neurons in fixed tissue. We found that brains subjected to Nup153 RNAi exhibited an almost complete absence of transfected cells within the IZ and CP regions unlike the control brains, consistent with a role for the Nup153 in recruitment of Kif1a to the NE to regulate neuronal migration (Supplemental Figure 6). Most of the Nup153 shRNA expressing cells were located in the SVZ and exhibited a multipolar morphology, suggesting a defect in the multipolar-to-bipolar neuronal transition. This effect was similar to that observed for Kif1a KD. However, in Kif1a knockdown brains, neighboring non-transfected cells were also arrested in the SVZ in a multipolar state. Transfected cells, and the neighboring cells also expressed Tbr1 – a marker normally expressed only by the postmitotic projection neurons^20, 21^. This non-cell autonomous effect of Kif1a knockdown was not phenocopied by Nup153 knockdown, evidenced by the unaltered Tbr1 band in the Nup153 shRNA expressing brains (Supplemental Figure 6).

The Kif1a tail region was originally used to pull down Nup153 from adult rat brain lysate. This region localizes to NE in nonneuronal cells, RGP cells and in postmitotic neurons, suggesting an important role for Nup153 in anchoring Kif1a to the NE. To test this directly, we used a commercially available Nup153 siRNA. Transfection of HeLa cells with this reagent reduced Nup153 expression as judged by immunostaining. Kif1a tail region expression in Nup153-depleted cells resulted in a dramatic loss of Kif1a NE localization (Figure 3). This result indicates that Nup153 is necessary for Kif1a NE localization.

**Figure 3.**
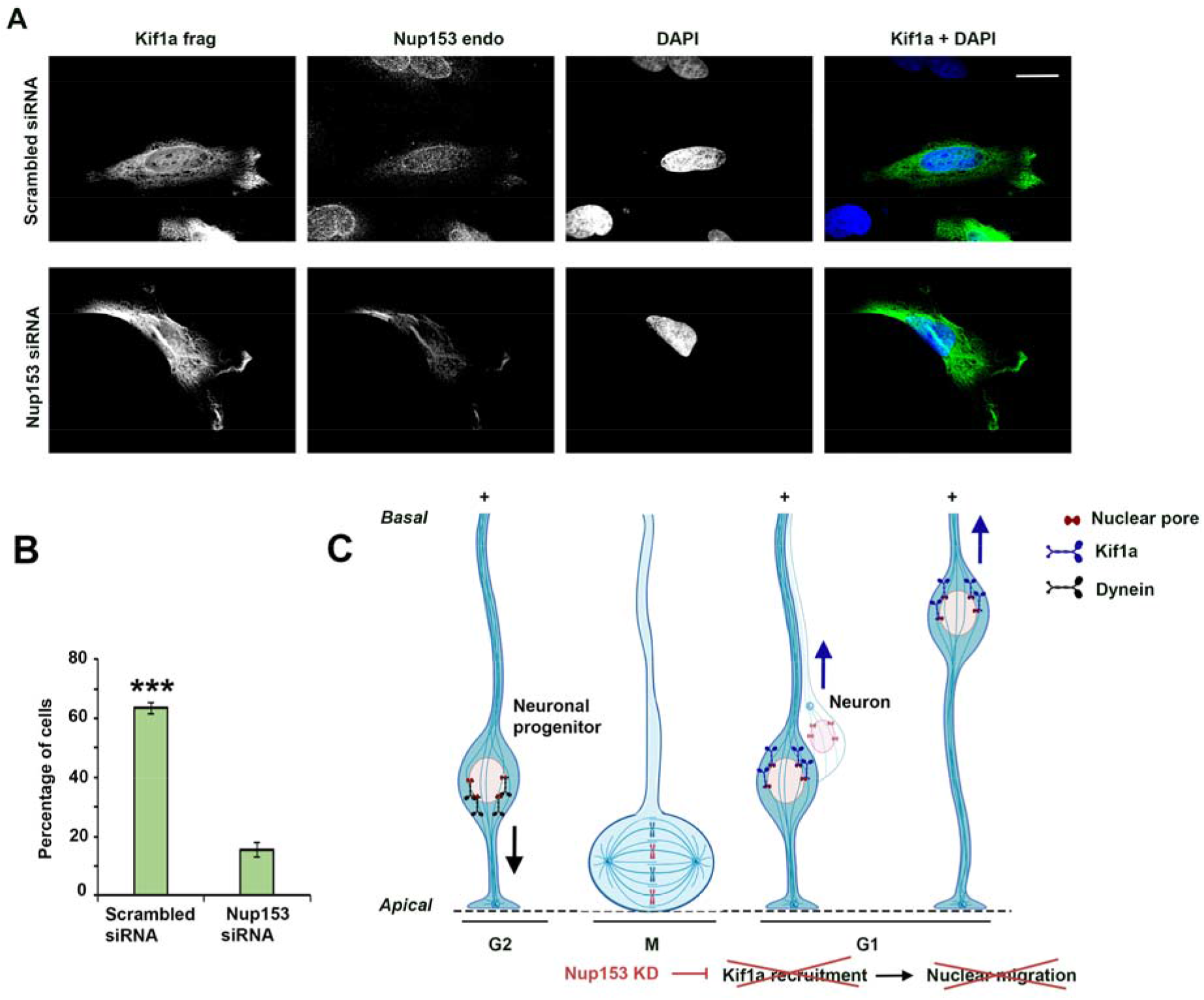
Nup153 is required for HeLa cell NE Kif1a recruitment. **A**. HeLa cells were transfected with either Nup153 siRNA or scrambled siRNA and fixed after 48 hours. 20 hours prior to fixation, the cells in both the groups were also transfected with EGFP tagged KIF1A tail fragment. Following fixation, the cells were stained with anti-Nup153 antibody as well as anti-EGFP antibody. Nup153 siRNA mediated loss of endogenous Nup153 (as determined by antibody staining) also led to the loss of NE decoration by KIF1A tail fragment. Scale bar 20 μm. **B**. Quantification of NE decoration by KIF1A tail fragment in the scrambled siRNA and Nup153 siRNA groups. Experiment was reproduced three independent times (over 50 cells per condition and per experiment were counted. _∗∗_p < 0.01; _∗∗∗_p < 0.001; error bars = SEM) **C**. Model for basal INM. Kif1a is shown to be recruited to RGP cell nuclear pores via Nup153, knockdown of which leads to the loss of Kif1a NE recruitment and inhibition of basal INM.

We previously found that INM is driven by kinesin- and dynein-mediated nuclear transport along an array of uniformly oriented microtubules^3^. During INM, the centrosome remains at the ventricular surface as the nucleus oscillates in basal and apical directions. This centrosome-independent nuclear migration suggested that the motor proteins may act from the nuclear surface, a relatively unusual mechanism in vertebrates. We now provide the first evidence that the kinesin, Kif1a is present at the nuclear surface and that RNAi for a gene responsible for Kif1a nuclear recruitment inhibits basal INM. A previous study demonstrated that HeLa cells depleted of Nup153 by RNAi can still carry out general nucleocytoplasmic transport^22^, suggesting that this is not the mechanism responsible for inhibition of INM in RGPs.

Unlike Kif1a RNAi, Nup153 RNAi resulted in defects in cell cycle progression in the RGP cells such that cells were arrested in G1 phase. The mechanism of how Nup153 facilitates G1/S transition has been identified in hepatocellular carcinoma (HCC) cells^23^. NUP153 negatively regulates the expression of P15^INK4b^ (cyclin-dependent kinase inhibitor 2B)^23^, which is known to trigger G1/S phase arrest by inactivating the Cyclin D-CDK4 /CDK6 complex^24, 25^. Our results indicating the role of Nup153 in regulating basal INM, which occurs during G1 imply that basal nuclear migration and G1/S transition are under the control of the same nucleoporin, which may ensure that these two processes always occur in proper sequence.

We previously found G2-specific mechanisms that recruit dynein to nuclear pores drive apical nuclear migration of brain progenitor cells^2^. Our present findings suggest a parallel mechanism where Kif1a is recruited to nuclear pores to drive basal nuclear migration.

## Abbreviations used in this paper

NE: Nuclear envelope
VZ: Ventricular Zone
IZ: intermediate zone
CP: Cortical plate
RGP: Radial glial progenitor
INM: interkinetic nuclear migration
KAND: KIF1A-associated neurological disorder

## REFERENCES

1. Taverna, E. & Huttner, W.B. Neural progenitor nuclei IN motion. Neuron 67, 906–914 (2010).

2. Hu, D.J., et al. Dynein recruitment to nuclear pores activates apical nuclear migration and mitotic entry in brain progenitor cells. Cell 154, 1300–1313 (2013).

3. Tsai, J.W., Lian, W.N., Kemal, S., Kriegstein, A.R. & Vallee, R.B. Kinesin 3 and cytoplasmic dynein mediate interkinetic nuclear migration in neural stem cells. Nat Neurosci 13, 1463–1471 (2010).

4. Del Bene, F., Wehman, A.M., Link, B.A. & Baier, H. Regulation of neurogenesis by interkinetic nuclear migration through an apical-basal notch gradient. Cell 134, 1055–1065 (2008).

5. Kosodo, Y. Interkinetic nuclear migration: beyond a hallmark of neurogenesis. Cell Mol Life Sci 69, 2727–2738 (2012).

6. Carabalona, A., Hu, D.J. & Vallee, R.B. KIF1A inhibition immortalizes brain stem cells but blocks BDNF-mediated neuronal migration. Nat Neurosci 19, 253–262 (2016).

7. Tsai, J.W., Chen, Y., Kriegstein, A.R. & Vallee, R.B. LIS1 RNA interference blocks neural stem cell division, morphogenesis, and motility at multiple stages. J Cell Biol 170, 935–945 (2005).

8. Tsai, J.W., Bremner, K.H. & Vallee, R.B. Dual subcellular roles for LIS1 and dynein in radial neuronal migration in live brain tissue. Nat Neurosci 10, 970–979 (2007).

9. Niwa, S., et al. Autoinhibition of a Neuronal Kinesin UNC-104/KIF1A Regulates the Size and Density of Synapses. Cell Rep 16, 2129–2141 (2016).

10. Chiba, K., Kita, T., Anazawa, Y. & Niwa, S. Insight into the regulation of axonal transport from the study of KIF1A-associated neurological disorder. J Cell Sci 136 (2023).

11. Boyle, L., et al. Genotype and defects in microtubule-based motility correlate with clinical severity in KIF1A-associated neurological disorder. HGG Adv 2 (2021).

12. Stucchi, R., et al. Regulation of KIF1A-Driven Dense Core Vesicle Transport: Ca(2+)/CaM Controls DCV Binding and Liprin-alpha/TANC2 Recruits DCVs to Postsynaptic Sites. Cell Rep 24, 685–700 (2018).

13. Duheron, V., Chatel, G., Sauder, U., Oliveri, V. & Fahrenkrog, B. Structural characterization of altered nucleoporin Nup153 expression in human cells by thin-section electron microscopy. Nucleus 5, 601–612 (2014).

14. Bastos, R., Lin, A., Enarson, M. & Burke, B. Targeting and function in mRNA export of nuclear pore complex protein Nup153. J Cell Biol 134, 1141–1156 (1996).

15. Enarson, P., Enarson, M., Bastos, R. & Burke, B. Amino-terminal sequences that direct nucleoporin nup153 to the inner surface of the nuclear envelope. Chromosoma 107, 228–236 (1998).

16. Fahrenkrog, B., et al. Domain-specific antibodies reveal multiple-site topology of Nup153 within the nuclear pore complex. J Struct Biol 140, 254–267 (2002).

17. Paulillo, S.M., et al. Nucleoporin domain topology is linked to the transport status of the nuclear pore complex. J Mol Biol 351, 784–798 (2005).

18. Paulillo, S.M., Powers, M.A., Ullman, K.S. & Fahrenkrog, B. Changes in nucleoporin domain topology in response to chemical effectors. J Mol Biol 363, 39–50 (2006).

19. Liu, J.S., et al. Molecular basis for specific regulation of neuronal kinesin-3 motors by doublecortin family proteins. Mol Cell 47, 707–721 (2012).

20. Andero, R., et al. Effect of 7,8-dihydroxyflavone, a small-molecule TrkB agonist, on emotional learning. Am J Psychiatry 168, 163–172 (2011).

21. Englund, C., et al. Pax6, Tbr2, and Tbr1 are expressed sequentially by radial glia, intermediate progenitor cells, and postmitotic neurons in developing neocortex. J Neurosci 25, 247–251 (2005).

22. Mackay, D.R., Elgort, S.W. & Ullman, K.S. The nucleoporin Nup153 has separable roles in both early mitotic progression and the resolution of mitosis. Mol Biol Cell 20, 1652–1660 (2009).

23. Gan, C., et al. NUP153 promotes HCC cells proliferation via c-Myc-mediated downregulation of P15(INK4b). Dig Liver Dis 54, 1706–1715 (2022).

24. Hannon, G.J. & Beach, D. p15INK4B is a potential effector of TGF-beta-induced cell cycle arrest. Nature 371, 257–261 (1994).

25. Yaginuma, Y., et al. Analysis of the Rb gene and cyclin-dependent kinase 4 inhibitor genes (p16INK4 and p15INK4B) in human ovarian carcinoma cell lines. Exp Cell Res 233, 233–239 (1997).

