## Supplemental figures and legends for "The Nucleoporin Nup153 Is the Anchor for Kif1a during Basal Nuclear migration in Brain Progenitor Cells"

**Supplemental Figure 1**

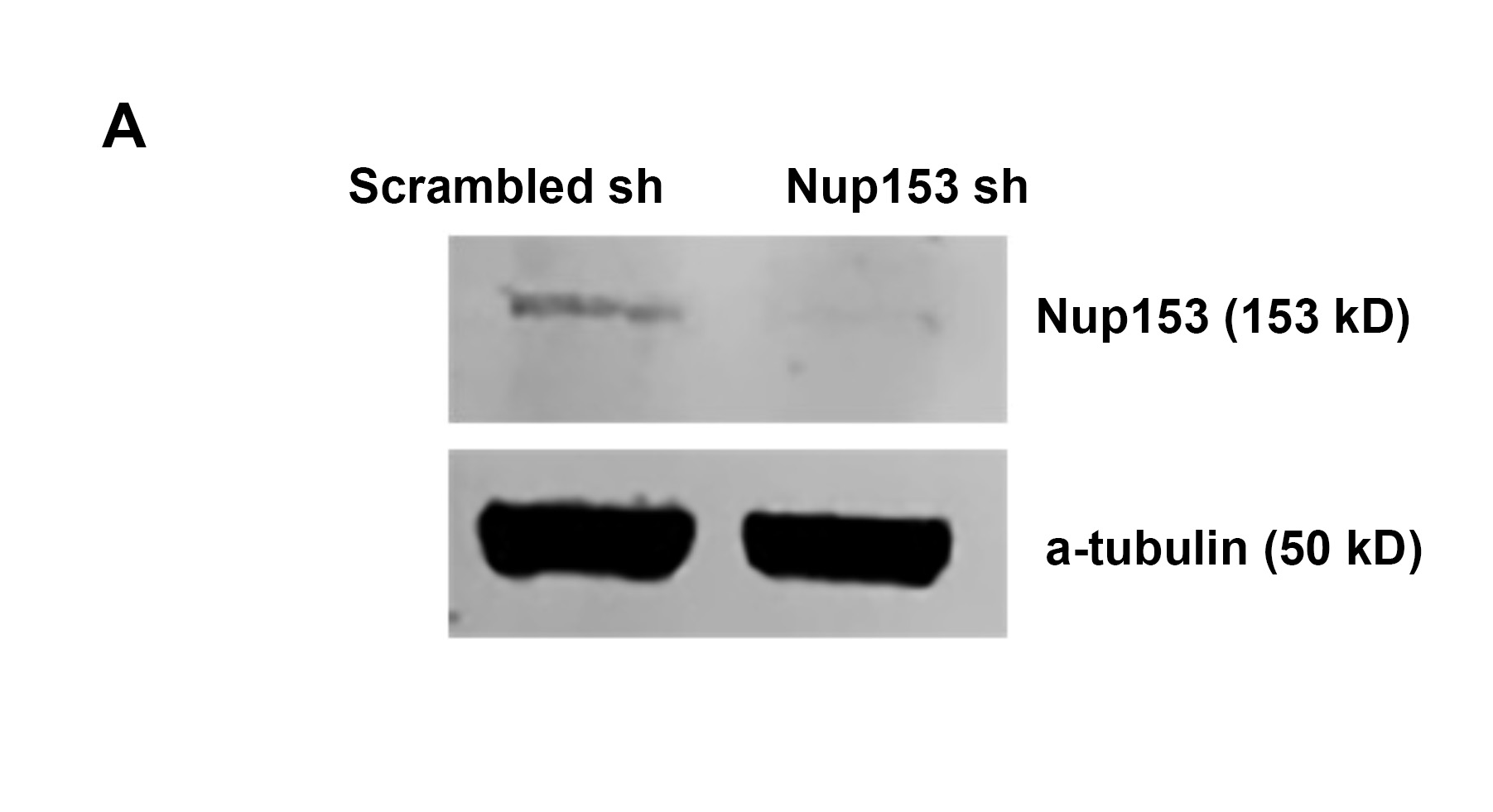

**SUPPLEMENTAL FIGURE 1:**

**Nup153 shRNA efficiently knocks down Nup153 protein expression in C6 glioma cells.**

**A.** C6 rat glioma cells were transfected with Nup153 shRNA or scrambled shRNA for 72 h. Western blot analysis confirms successful protein KD.

**Supplemental Figure 2**

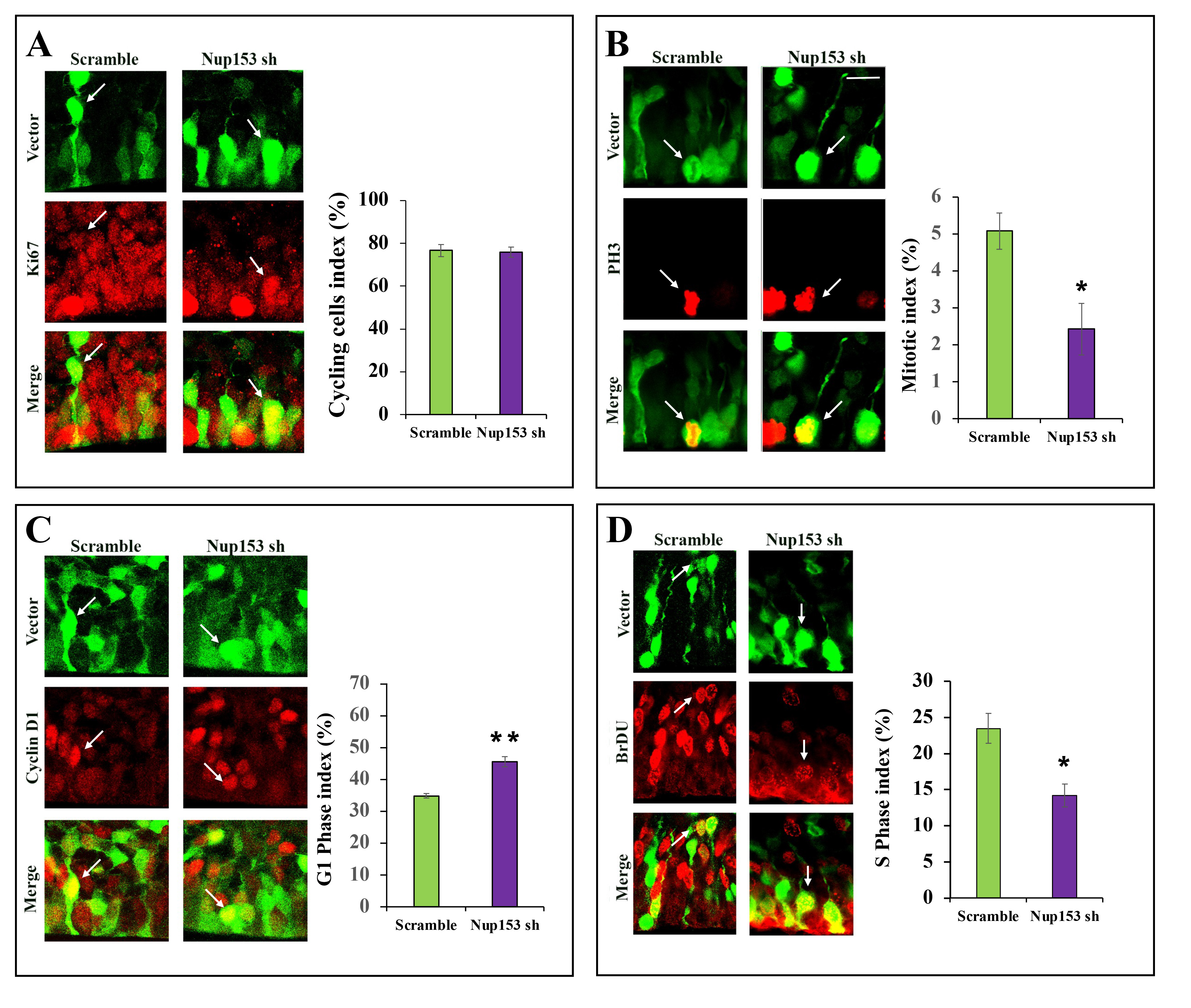

**Nup153 RNAi in RGP cells leads to an increase in G1 phase cells.**

**A-D.** E16 embryonic brains were in utero electroporated with either scrambled shRNA or shRNA for Nup153. Brains were then fixed at E19 and stained with cell cycle markers as follows: Ki67 (cycling cells), Cyclin D1 (G1 phase), phospho-histone H3 (PH3) (late G2/M phases). We also investigated the effect of Nup153 RNAi on S phase by BrDU pulse labeling. A comparable percentage of control and Nup153 knockdown RGP cells expressed Ki67 (scramble 76.7 ± 2.7%; n=3, Nup153 shRNA 75.9 ± 2.4%; p=0.8; n=3), suggesting that Nup153 has no gross effect on the fraction of cycling cells, although a subpopulation in each case escaped the cell cycle. There was a significant increase in the percentage of RGP cells positive for cyclin D1 (scramble 35.9 ± 0.8%; n=3, Nup153 shRNA 45.6 ± 1.6%; p=0.002; n=3) and a corresponding decrease in the percentage of RGP cells positive for phospho-histone H3 (scramble 5.1 ± 0.5%; n=3, Nup153 shRNA 2.4 ± 0.7%; p=0.04; n=3) as well as BrDU (scramble 23.5 ± 2.1%; n=3, Nup153 shRNA 14.2 ± 1.6%; p=0.02; n=4. ∗∗p < 0.01; ∗∗∗p < 0.001; error bars = SEM). Scale bar 15 μm.

**Supplemental Figure 3**

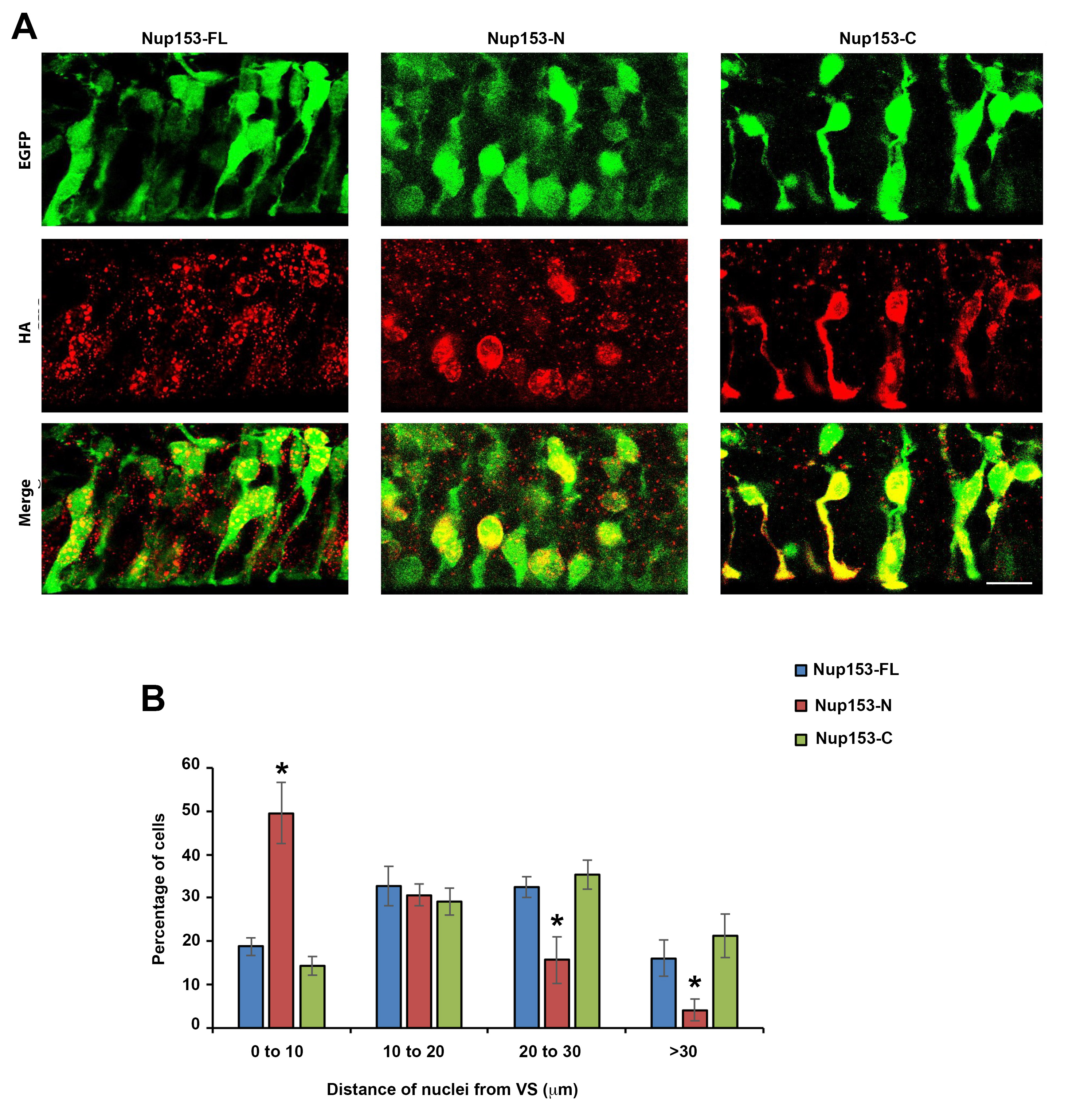

**Mechanistic insight from the Nup153 functional domains.**

**A,B.** E16 brains were in utero electroporated with the full-length, or the N-terminal domain or the C-terminal domain of Nup153, with HA tag fused to its N-terminus and EGFP tag that is cytoplasmic. Analysis was performed at 3 d.p.i.

**A.** Fixed images of the VZ from the electroporated brains stained for EGFP to enhance the signal and for HA to determine the localization of each cDNA construct. Scale bar 15 μm.

**B.** Quantification of the distance between RGP nuclei and the VS across the various conditions. Expression of full-length Nup153 or the Nup153-C domain alone had no detectable effect on the position of the nucleus within the RGP cell However, Nup153-N domain expression in RGP cells caused a pronounced accumulation of most transfected cells with somata located close to the ventricular surface (0 – 10 μm: Nup153-FL, 18.7 ± 2.1%, n = 3; Nup153-N, 49.6 ± 7.2%, n = 4, p = 0.02; Nup153-C, 14.3 ± 2.1%, n = 3; p = 0.2; 10 – 20 μm: Nup153-FL, 32.7 ± 4.5%, n = 3; Nup153-N, 30.7 ± 2.6%, n = 4, p = 0.65; Nup153-C, 29.11 ± 3.2%, n = 3, p = 0.6; 20 – 30 μm: Nup153-FL, 32.6 ± 2.4%, n = 3; Nup153-N, 15.7 ±4.3%, n = 4, p = 0.06; Nup153-C, 35.3 ± 3.3%, n = 3, p = 0.5; >30 μm: Nup153-FL, 16.0 ± 4.2%, n = 3; Nup153-N, 4.1 ± 2.5%, n = 4, p = 0.03; Nup153-C, 21.3 ± 5.1%, n = 3, p = 0.5; ∗∗p < 0.01; ∗∗∗p < 0.001; error bars = SEM).

**Supplemental Figure 4**

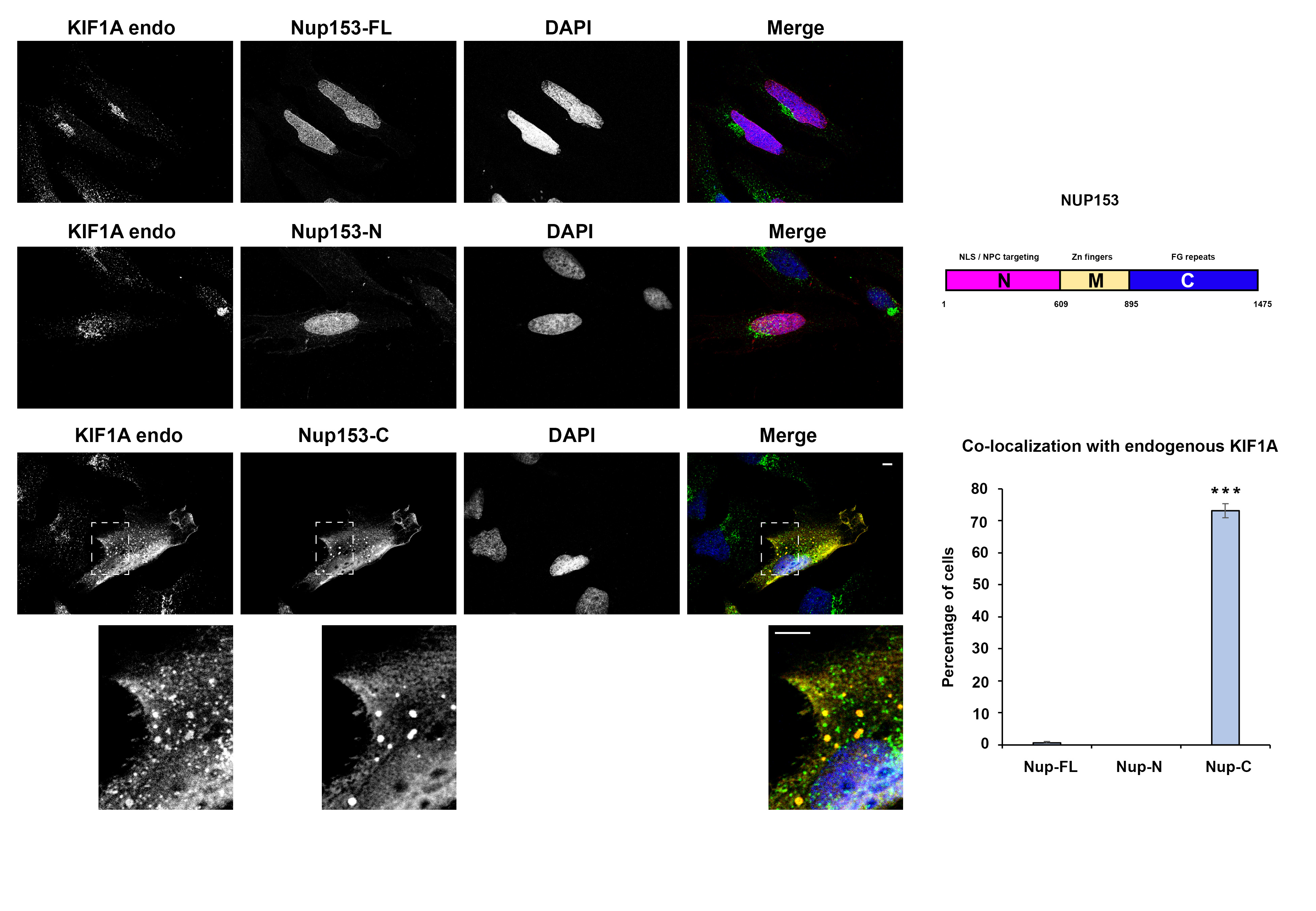

**The C-terminal domain of Nup153 colocalizes with the endogenous Kif1a.**

**A.** HeLa cells expressing full-length, or the N-terminal domain or the C-terminal domain of Nup153 with mCherry-tag that is fused at its N-terminal are stained for endogenous Kif1a. Area in the white dotted boxes is enlarged below. Endogenous KIF1 shows clear colocalization with Nup153-C fragment. Scale bars 5 μm.

**B.** Schematic representation of Nup153 molecule.

**C.** Quantification of co-localization between Kif1a tail fragment and Nup153 fragments. (∗∗p < 0.01; ∗∗∗p < 0.001; error bars = SEM).

**Supplemental Figure 5**

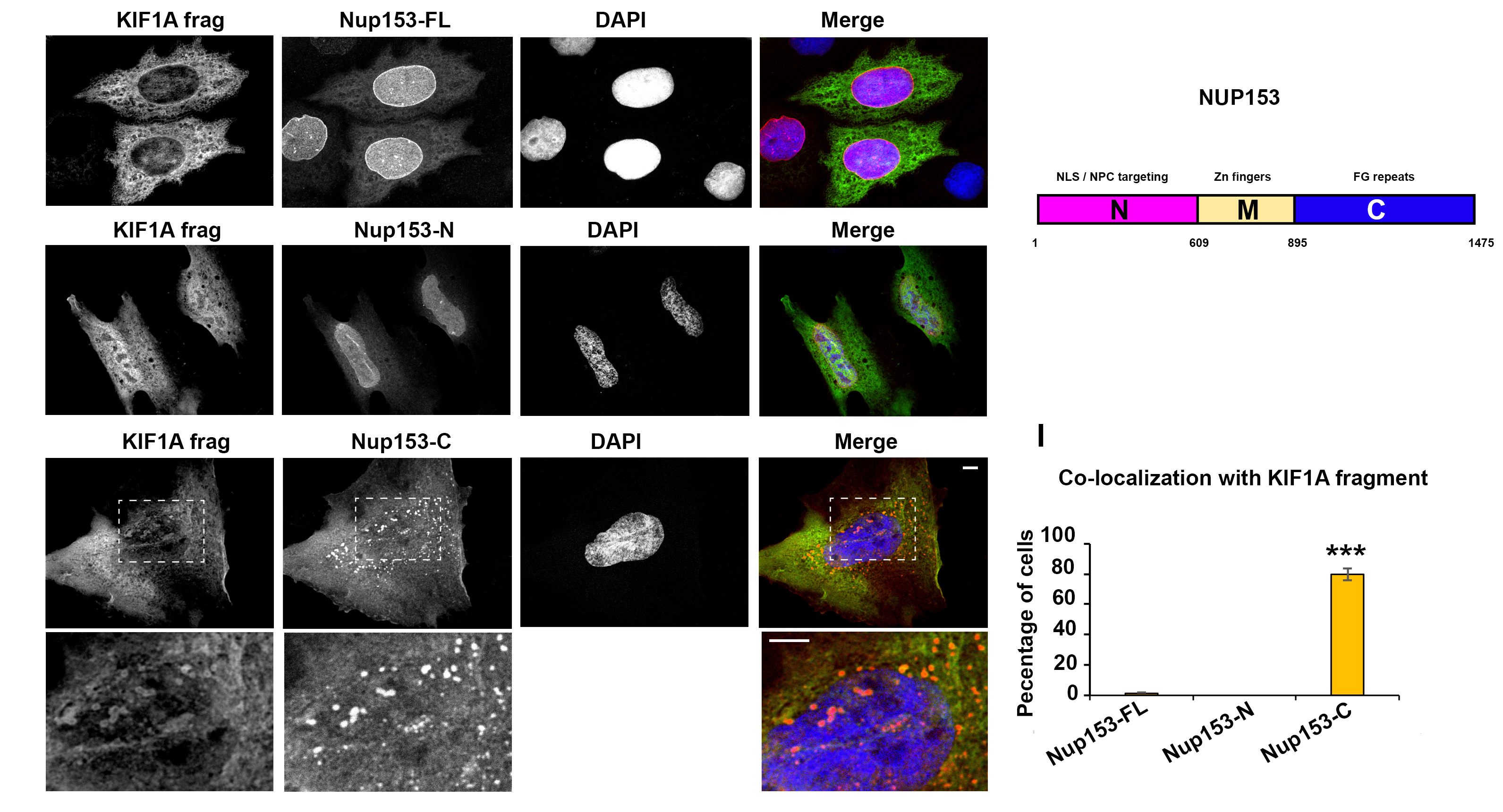

**The C-terminal domain of Nup153 colocalizes with the recombinant Kif1a.**

**A.** HeLa cells co-expressing KIF1A tail fragment with EGFP-tag that is fused at its N-terminal and full-length, or the N-terminal domain or the C-terminal domain of Nup153 with mCherry-tag that is fused at its N-terminal. Area in the white dotted boxes is enlarged below. Endogenous KIF1 shows clear colocalization with Nup153-C fragment. Scale bars 5 μm.

**B.** Schematic representation of Nup153 molecule.

**C.** Quantification of co-localization between Kif1a tail fragment and Nup153 fragments. (∗∗p < 0.01; ∗∗∗p < 0.001; error bars = SEM).

**Supplemental Figure 6**

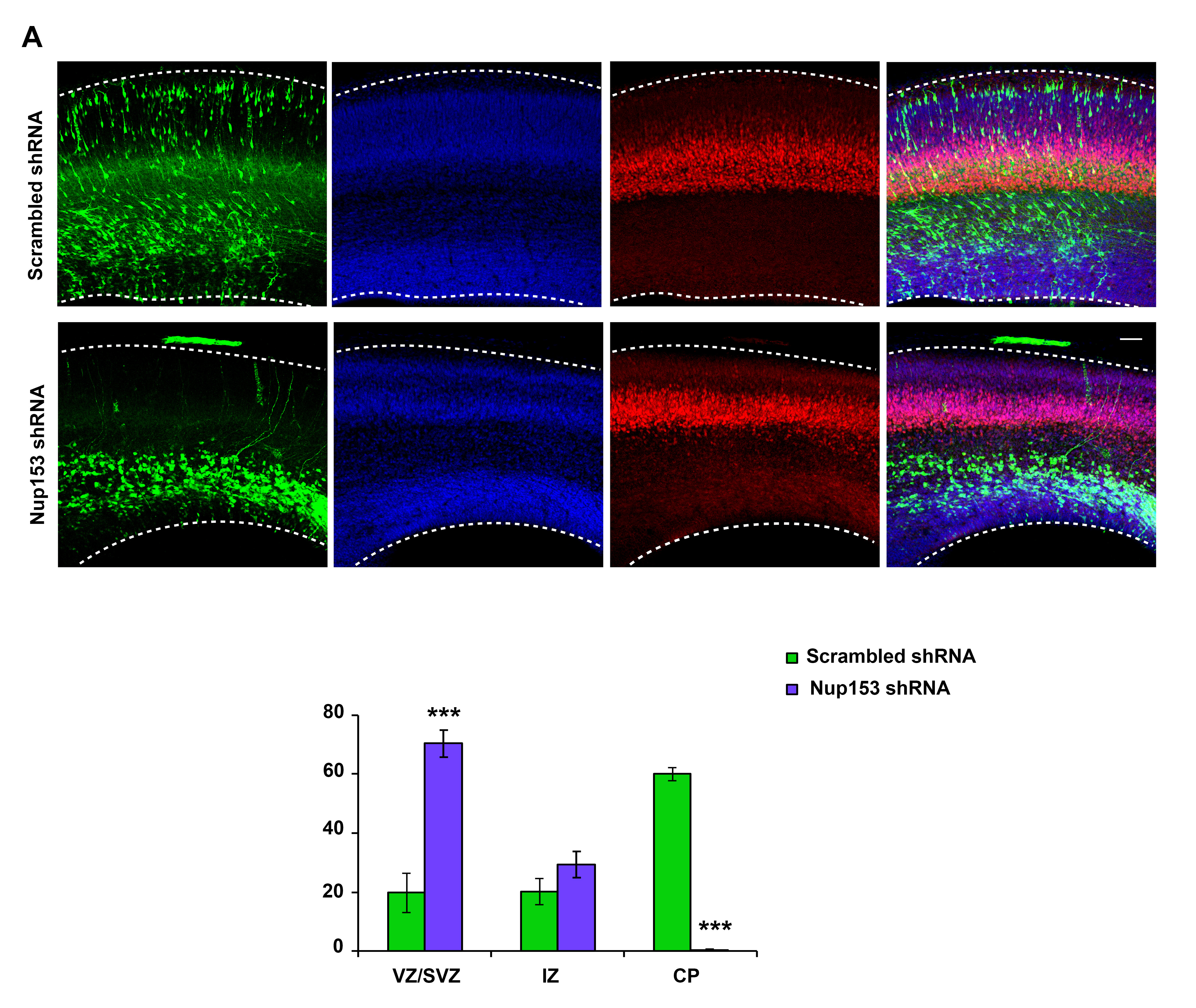

**Role for Nup153 in post-mitotic neuronal migration.**

E16 rat embryonic brains were in utero electroporated with shRNA for Nup153 or scrambled shRNA,. Analysis was performed 4 d.p.i.

**A.** Images of the neocortex injected with scrambled or Nup153 shRNA and stained for Tbr1. Scale bar 50 μm.

**B.** Quantification of the proportion of transfected cells in each layer of the neocortex across conditions. Depletion of Nup153 leads to an accumulation of the majority of transfected neurons at the multipolar stage within the SVZ/lower IZ but does not affect Tbr1 expression in nearby neurons. (VZ/SVZ: scramble shRNA, 19.8 ± 6.6%, n = 3; Nup153 shRNA, 70.3 ± 4.6%, n = 3; p = 0.0004; IZ: scramble shRNA, 20.2 ± 4.3%, n = 3; Nup153 shRNA, 29.2 ± 4.4%, n = 3, p = 0.07; CP: scramble shRNA, 60.0 ± 2.3%, n = 3; Nup153 shRNA, 0.4 ± 0.2%, n = 3, p = 0.0001; ∗∗p < 0.01; ∗∗∗p < 0.001; error bars = SD).

**Materials and Methods**

**Ethics statement**

All the experiments were done in accordance with the animal welfare guidelines and the guidance of the Institutional Animal Care and Use Committee at Columbia University (New York, NY).

**In utero electroporation**

Plasmids encoding for shRNAs or cDNA were injected into the lateral ventricles of embryonic rat brain at E16 and electroporated as described^1^. Specifically, timed pregnant Sprague Dawley E16 rats were anaesthetized with a mixture of ketamine/xylazine (respectively at 90 and 5mg/kg), administered intraperitoneally. To avoid excessive heat-loss during the surgical procedure, an external heating source was provided. For pain management, bupivacaine (2mg/kg) was administered via a subcutaneous injection at the site of the future incision, and buprenorphine (0.05mg/kg) was administered by a subcutaneous injection. This dose of buprenorphine was re-administered to the animal every 8–12 hours, for up to 48 hours following the surgery. Abdominal cavity was opened, and uterine horns were exposed and trans-illuminated for clear identification of the brain ventricles. For easy visualization of the DNA in the brain ventricular space, a nontoxic dye (Sigma, F7252) was added to the DNA before surgery and injected with a sharpened glass needle. After injection, plasmids were further electroporated by discharging a 4000 mF capacitor charged to 50V with a BTX ECM 830 electroporator (BTX Harvard Apparatus, Holliston, MA, USA). The voltage was discharged in five electrical pulses at 950ms intervals via 7mm electrodes placed on the head of the embryo across the uterine wall. The embryos were returned to the abdominal cavity, and the wound was closed. Rats were monitored every day after surgery.

**Immunohistochemistry and live imaging**

For live imaging, brains were harvested 3 days after the electroporation. The dissected rat brains were embedded in 4% low-melting agarose (IBI Scientific, IB70057) prepared in artificial cerebrospinal fluid^1^ and sliced coronally (300-µm) on a vibratome (Leica microsystems). The slices were placed on 0.4μm, 30mm diameter Millicell-CM inserts (Millipore) in cortical culture medium containing 25% HBSS (Life Technologies, 24020-117), 47% basal MEM (Life Technologies, 21010-046), 25% normal horse serum (Life Technologies, 26050-088), 1% penicillin/streptomycin/glutamine (Life Technologies, 10378-016), and 2% of 30% glucose (Sigma, G5767) in a 50-mm glass-bottom dish (MatTek Corporation, P50G-0-14-F) and imaged on an IX80 laser scanning confocal microscope (Olympus FV1000 Spectral Confocal System) at intervals of 10 min for up to 24 h.

For fixed imaging, embryonic (E19 or E20) rat brains were harvested and fixed with chilled saline and 4% paraformaldehyde (PFA; EMS, wt/vol) and then incubated in 4% PFA overnight. Brains were then embedded in 4% of agarose (Sigma, A9539) and sliced sectioned coronally (100μm) on a vibratome (Leica microsystems). After blocking in 5% normal donkey serum (Sigma, D9663) in PBS-Triton 0.5% for 1 h, slices were incubated with primary antibodies diluted in the blocking solution, overnight on a shaker, at 4°C. Secondary antibodies (1:500) and DAPI (Thermo Scientific, 62248; 1:10,000 dilution) were diluted in PBS and incubated for 2 h at room temperature. Slices were mounted with Aqua-Poly mounting media (Polysciences, 18606).

Antibodies used in this study were: Tbr2 (Millipore, AB2283), KI67 (Millipore, AB9260), Cyclin D1 (ThermoScientific, RM-9104-S0), phospho-histone H3 (Abcam, ab14955), BrDU (Abcam, ab6326), Kif1a (Abcam, ab240222), HA (Sigma-Aldrich, H6908), mCherry (Abcam, ab167453), GFP (Abcam, ab1218) and donkey fluorophore–conjugated secondary antibodies (Jackson Labs). Nup153 antibody was a kind gift from Dr. Brian E. Burke, A*STAR Skin Research Labs (A*SRL) – Biopolis, Singapore.

**RNAi and constructs**

shRNA expressing constructs were designed to target internal gene sequences unique to rat Nup160 and rat Nup153, in a pRetro-U6G vector (Cellogenetics), which also expressed soluble GFP to label-transfected cells. The shRNA sequence for Nup160 was 5′-CCCTATGTGAATCTGCATAAT-3′, and the sequence for Nup153 was 5′-ACTTCAGTTTCTGGTCGCAAGATAA-3′. Scrambled Nup153 shRNA sequence was 5’-GTGCATAATATATGCGATCTATGCC-3’. Full-length human Nup153 and Nup153-C as well as Nup153-N were obtained from Addgene (Addgene, plasmid no.s #64268, #64318, #64317) and cloned into pCAGIG vector (Addgene, plasmid no. 11159). Full-length and functional domain constructs of Nup153 were HA (YPYPVPDYA). Human kif1a-tail fragment 650–1105 aa^2^ was a gift from Dr. Casper Hoogenraad (Genentech).

**Western blot**

shRNAs and were transfected in rat C6 brain glioma cells cultured in DMEM supplemented with 10% FBS and 1% penicillin/streptomycin and maintained at 37°C with 5% CO_2_. Transfection of cultured cells with shRNAs for Nup153 and Nup153 scrmabled was performed using a Lonza Nucleofector kit V and an Amaxa Nucleofector, according to the manufacturer’s instructions. Cells transfected with shRNAs were collected 72 h (for shRNAs) after transfection.

Cells transfected with shRNAs and brain samples were lysed on ice in Lysis Buffer (pH 7.2, 50 mM Tris-HCl, 150 mM NaCl, 1% Triton X-100, and 0.5% deoxycholic acid buffer) containing 1 mM DTT and a protease inhibitor cocktail (Sigma, P8340). Purified lysates were loaded on a polyacrylamide gel and transferred to a polyvinylidene difluoride membrane. The membrane was blocked in PBS with 0.5% powdered milk, incubated with primary antibodies diluted in PBS with 0.1% of powder milk, and washed and incubated with secondary LI-COR antibodies in PBS. Imaging of the blots was performed using an Odyssey system (LI-COR). Antibody from Abcam (ab 24700) was used to detect Nup153 on the Western blot.

**Imaging and statistical analysis**

All images were collected with an IX80 laser scanning confocal microscope (Olympus FV1000 Spectral Confocal System). Brain sections were imaged using a 60× 1.42 NA oil objective or a 10× 0.40 NA air objective. All images were analyzed using ImageJ software (National Institutes of Health).

All statistical analysis was performed using Prism (GraphPad Software). Unpaired *t* test (two-tailed) was used to determine significance between two groups. Definition of statistical significance was P < 0.05.

For each experimental condition, at least 3 embryos were collected from at least three different mothers.

**References:**

1.	Baffet, A.D., Hu, D.J. & Vallee, R.B. Cdk1 Activates Pre-mitotic Nuclear Envelope Dynein Recruitment and Apical Nuclear Migration in Neural Stem Cells. *Dev Cell* **33**, 703-716 (2015).

2.	Stucchi, R.*, et al.* Regulation of KIF1A-Driven Dense Core Vesicle Transport: Ca(2+)/CaM Controls DCV Binding and Liprin-alpha/TANC2 Recruits DCVs to Postsynaptic Sites. *Cell Rep* **24**, 685-700 (2018).
